# Genomic copy-number loss is rescued by self-limiting production of DNA circles

**DOI:** 10.1101/255471

**Authors:** Andrés Mansisidor, Temistocles Molinar, Priyanka Srivastava, Hannah Blitzblau, Hannah Klein, Andreas Hochwagen

## Abstract

Copy-number changes generate phenotypic variability in health and disease. Whether organisms protect against copy-number changes is largely unknown. Here, we show that *Saccharomyces cerevisiae* monitors the copy number of its ribosomal DNA (rDNA) and rapidly responds to copy-number loss with the clonal amplification of extrachromosomal rDNA circles (ERCs) from chromosomal repeats. ERC production is proportional to repeat loss and reaches a dynamic steady state that responds to the addition of exogenous rDNA copies. ERC levels are also modulated by RNAPI activity and diet, suggesting that rDNA copy number is calibrated against the cellular demand for rRNA. Lastly, we show that ERCs reinsert into the genome in a dosage-dependent manner, indicating that they provide a reservoir for ultimately increasing rDNA array length. Our results reveal a DNA-based mechanism for rapidly restoring copy number in response to catastrophic gene loss that shares fundamental features with unscheduled copy-number amplifications in cancer cells.

## Introduction

Repetitive gene arrays, like other repeated DNA sequences, are hotspots of genome plasticity due to their susceptibility to non-allelic homologous recombination (NAHR). One of the best-studied examples is the ribosomal RNA gene cluster (rDNA), which codes for the RNA components of the ribosome and is arranged in one or multiple gene arrays in most eukaryotes (Eickbush and Eickbush, 2007). In the human population, rDNA copy number varies frequently and widely (Gibbons et al., 2014; Stults et al., 2008), likely because only a fraction of rRNA genes is highly transcribed (Dammann et al., 1995). Low rDNA copy numbers, however, are linked to increased DNA damage sensitivity and cancer (Ide et al., 2010; Wang and Lemos, 2017; Xu et al., 2017).

Perhaps as a result of these selective pressures, several forms of rDNA copy number regulation have been observed. The single rDNA array of the yeast *Saccharomyces cerevisae* gradually shrinks and grows to restore array size after large-scale copy-number changes (Kobayashi et al., 1998). Relative copy number regulation is also implied for the human 5S and 45S rRNA genes, which co-vary in copy number despite mapping to unlinked repetitive arrays (Gibbons et al., 2015). In addition, rDNA copy number in yeast and flies is coupled with metabolic demands through trans-generational array contraction and repeat amplification (Aldrich and Maggert, 2015; Ha and Huh, 2011; Jack et al., 2015). Most of these changes, however, occur over the course of hundreds of cell divisions. Thus, it is unclear if copy number control is purely arising from selective pressures that differentially expand cell populations with advantageous copy number, or if cells also directly assess the number of rDNA copies in the short term.

Here, we tested this question by creating a series of large-scale deletions in the rDNA of *S. cerevisiae*. We show that rDNA copy number loss triggers the rapid and proportional production of extrachromosomal rDNA circles (ERCs). ERC production is self-limiting and responds to diet and to rRNA transcriptional output. Strains with higher ERC levels exhibit more frequent ERC re-integrations into the chromosomal rDNA array, providing a mechanism for array expansion.

## Results

### ERC production responds to chromosomal rDNA copy number

To probe the cellular response to copy-number changes, we created a collection of strains with different rDNA array lengths in *S. cerevisiae*. Partial deletions of the single yeast rDNA array were induced by ends-apart insertion of a *URA3*-marked targeting plasmid (Figure 1A). Based on pulsed-field gel electrophoresis (PFGE) analysis, we selected 12 strains with rDNA arrays comprising between ~90 and ~180 repeats (Figures 1B, S1A, Table S1). These array lengths roughly span the copy number considered wild-type for this organism (100-200 repeats) (Salim et al., 2017; Schweizer et al., 1969). Array lengths were stable at the population level over at least 150 divisions even though the rDNA fork barrier protein Fob1, which promotes expansion of critically short arrays (Kobayashi et al., 1998), was left intact. Minor expansion of short arrays was observed after repeated bottlenecking (Figure S1B), consistent with previous findings (Kobayashi et al., 1998).

**Figure 1:**
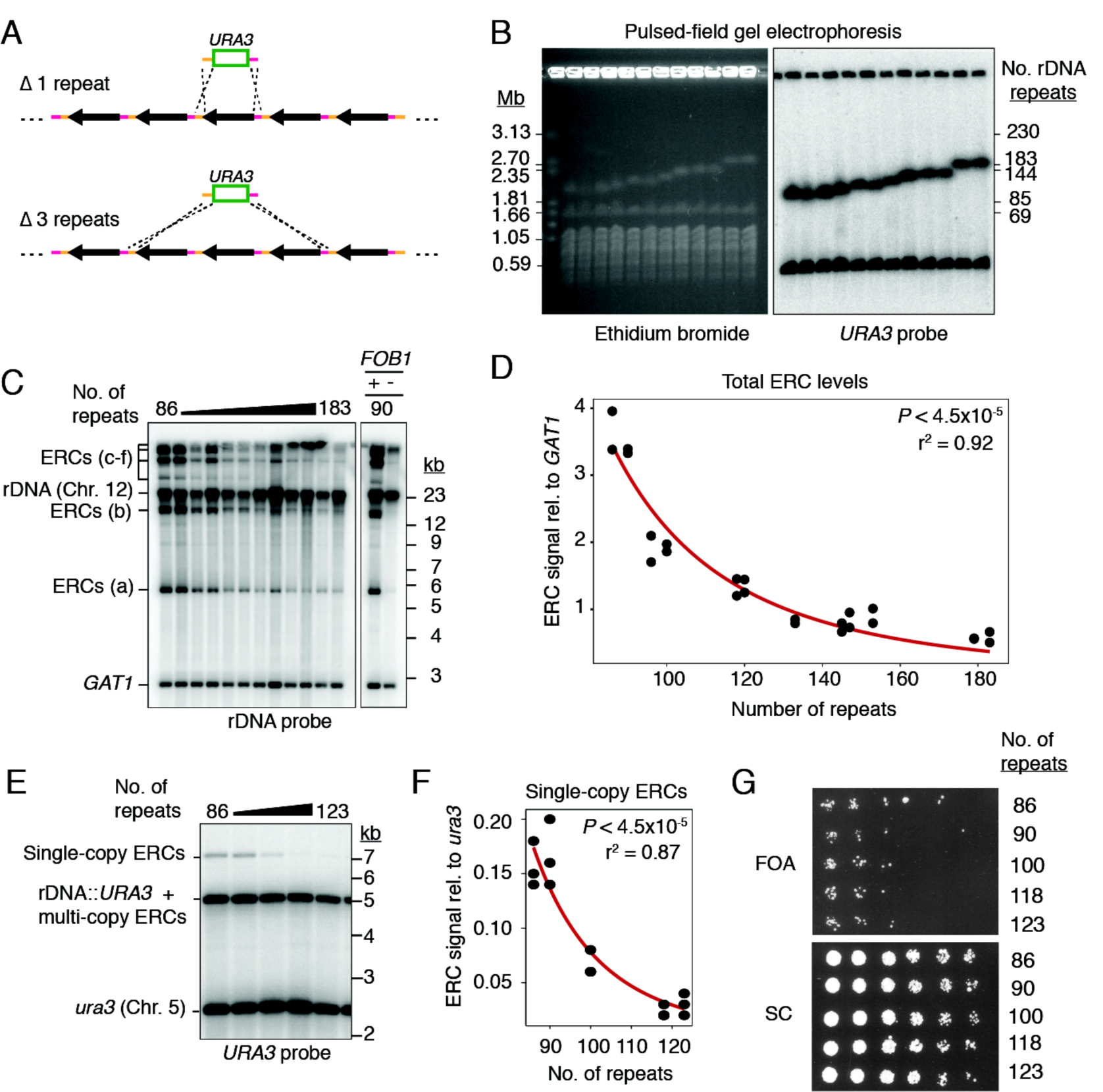
ERC levels anti-correlate with chromosomal rDNA array size. **(A)** Random deletion of chromosomal rDNA repeats using a *URA3*-targeting plasmid containing homologies to the rDNA intergenic sequence (orange and pink). **(B)** PFGE analysis of deletion strains. Southern probe (*URA3*) detects insertion in rDNA and *ura3* locus (Chr. 5). **(C)** Southern analysis of ERCs probing against rDNA sequence and *GAT1* (loading control). ERC signals labeled “a-f” based on electrophoretic mobility. **(D)** Total ERC signal (bands “a-e”) relative to *GAT1* as function of array size; *P* < 4.5x10^-5^ (Spearman’s rank correlation) and r^2^ fit to a power law function. **(E-F)** Southern analysis of *URA3*-marked ERCs as function of array size. **(G)** 3.5-fold dilutions of rDNA deletion strains spotted on synthetic complete (SC) media with and without 5-FOA.

Intriguingly, Southern assays revealed increased levels of ERCs in strains with short rDNA arrays. ERCs are generated by *FOB1*-stimulated intrachromatid NAHR (Defossez et al., 1999) and typically contain 1 to 5 rDNA repeats (Burkhalter and Sogo, 2004). Because of their circular nature, ERCs are detected above and below the sheared linear rDNA signal when undigested rDNA is separated by standard gel electrophoresis (Figure 1C) (Sinclair and Guarente, 1997). Quantification against a chromosomal loading control revealed a strong dependence on array size, with ~90-repeat strains exhibiting 5-10 times more ERCs than ~180-repeat strains (Figure 1D). This effect requires *FOB1* (Figure 1C) and is also apparent for ERCs originating from the rDNA repeat containing the *URA3* construct (Figure 1E). ERC signals are resistant to RecBCD exonuclease (Figure S1C-D) and exhibit the expected dependence on the repair factors *RAD52* and *RAD50*, but not *RAD51* (Figure S1E) (Park et al., 1999). We conclude that shortening of the rDNA array is linked to a proportional *FOB1*-dependent increase in ERC levels. This increase is not connected to the well-established accumulation of ERCs in replicatively old yeast cells (Sinclair and Guarente, 1997), because replicatively old cells (>15 divisions) comprise 0.003% of the logarithmically growing cultures used in this study, and lifespan is unaffected by a reduction of array size to 80 repeats (Saka et al., 2013).

### ERC formation is functionally separable from rDNA repeat instability

To test if high ERC levels reflect increased rDNA instability, we monitored the loss of the *URA3* construct. Cells were grown for 150-200 generations without selection before plating on 5-fluoroorotic acid (5-FOA), which kills cells expressing *URA3*. Irrespective of array length, approximately 1 in 4.0x10^4^ cells were able to grow on 5-FOA (Figure 1F). 5-FOA resistance was not the result of gene silencing because no recovery was observed after transfer to medium lacking uracil (Figure S1F) and loss of the *URA3* marker was evident by Southern analysis (Figure S1G). The fraction of FOA^R^ cells aligns with previous measurements of marker loss in wild-type cells (Defossez et al., 1999; Gangloff et al., 1996; Ganley and Kobayashi, 2011) and indicates that shortening the rDNA array down to ~90 repeats does not detectably increase rDNA instability.

The accumulation of *URA3*-marked ERCs without substantial marker loss from the array suggests that ERCs are copied from a chromosomal repeat. This notion is supported by the relative abundance of *URA3*-marked ERCs. Southern analysis after linearizing repeats showed that single-copy ERCs containing the *URA3* construct accumulate to ~15% of a genomic *ura3* loading control in ~90-repeat strains (Figure 1E). Isolation of RecBCD-resistant species further indicated that multi-copy ERCs derived from the same repeat are present at 2.5-3 times the level of single-copy ERCs (Figure S1H-I). Thus, ERC levels derived from single chromosomal repeats are orders of magnitude higher than the loss rates of the same repeats, strongly implying that ERCs are copied from array-encoded repeats.

### ERC production is rapid and self-limiting

To monitor the kinetics of ERC accumulation, we rendered ERC formation conditional by placing *FOB1* under the control of an estrogen-inducible expression system in a ~90-repeat strain (Figure 2A-B) (Benjamin et al., 2003). This system induces *FOB1* at near endogenous levels (Figure S2A). Cells maintained in log phase prior to *FOB1* induction recapitulated the low ERC levels of *fob1*Δ mutants (Figure 2B). Upon estradiol addition, all major ERC species became detectable within 6 divisions, with levels reaching a steady state after about 12 divisions. Analysis of single-copy ERCs derived from the *URA3*-marked repeats yielded similar kinetics and showed that ERC levels stabilized at ~20% relative to a genomic loading control (Figure S2B-C). This level is similar to the ERC levels observed in strains that have undergone 150-200 divisions (Figure 1E), suggesting that ERC levels reach a steady state that can be maintained for more than 150 divisions.

**Figure 2:**
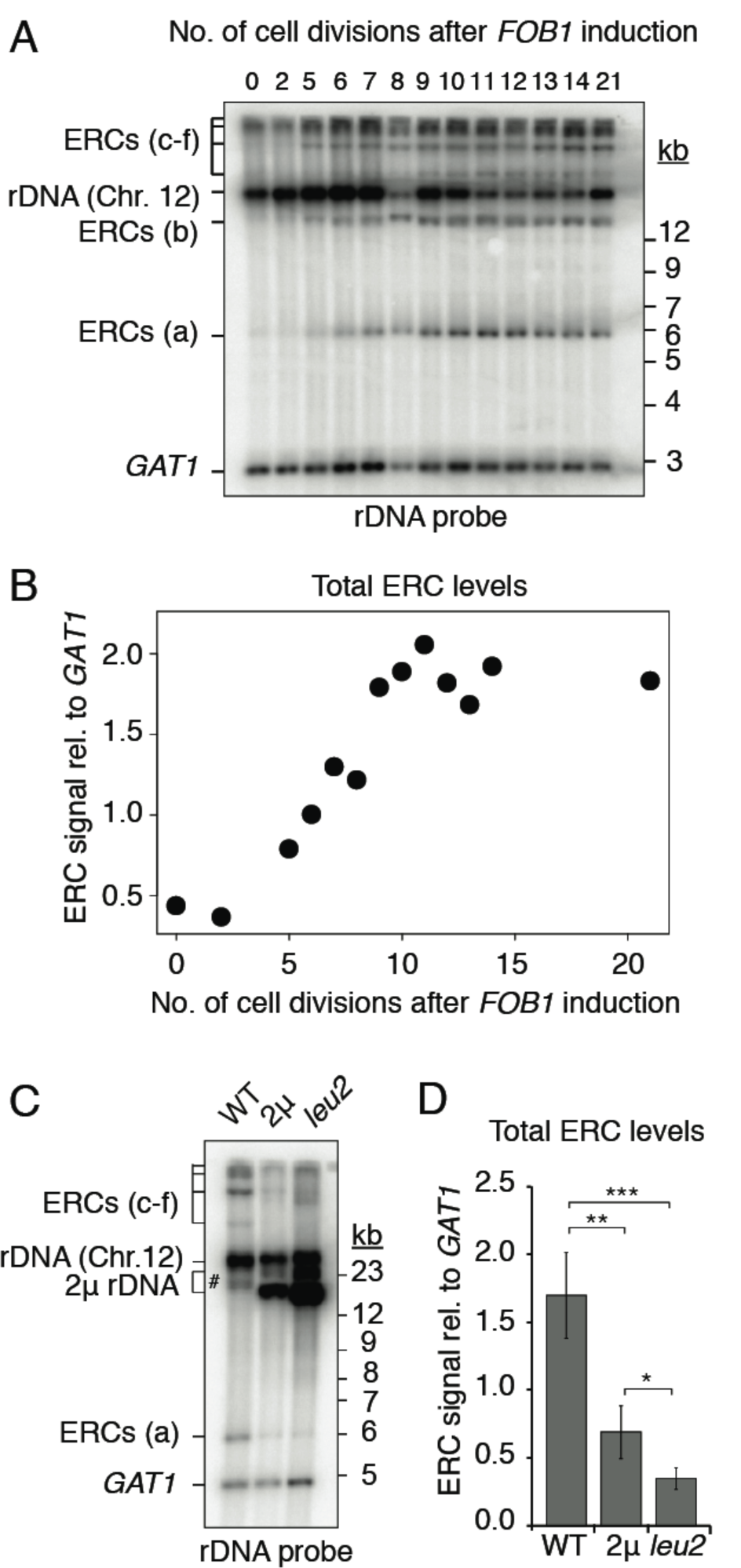
ERC accumulation responds to copy number. **(A-D)** Southern analysis of ERCs in ~90-repeat strains. **(A-B)** Kinetics of ERC accumulation (bands a-e) relative to *GAT1* after *FOB1* induction with 100nM beta-estradiol. **(C-D)** ERC levels (bands a, c, d, e) relative to *GAT1* after introduction of high-copy (2μ) or extra-high-copy (*leu2*) rDNA plasmids; ERC b (#) overlapped plasmid signal and was not quantified. \*\**P* < 0.01, \*\*\**P* < 0.001 (one-tailed *t*-test with Holm correction, n=4).

The rapid implementation of a steady-state ERC level that depends on array size implies that cells assess the total copy number of the rDNA and reduce ERC formation once sufficient rDNA copies have accumulated. To test this model, we transformed cells with an rDNA-bearing, high-copy plasmid whose copy number can be further amplified by selection in the absence of leucine (Figure 2C) (Chernoff et al., 1994). Transformation of this plasmid into a ~90-repeat strain led to a strong reduction in endogenous ERC levels and a further reduction when plasmid copy number was boosted in the absence of leucine (Figure 2C-D). The new steady-state levels demonstrate that cells assess total rDNA copy number. This experiment also indicates that copy-number is determined regardless of whether rDNA copies are chromosomal or episomal, implying negative-feedback regulation of ERC production.

### Diet and rRNA transcription modulate ERC levels

We sought to investigate the mechanisms by which cells monitor total rDNA copy number. Metabolic demand has long-term effects on rDNA copy number in yeast and flies, suggesting that cells respond to the need for rRNA (Aldrich and Maggert, 2015; Jack et al., 2015). To test if metabolic demand impacts ERC levels, we analyzed the effect of slower logarithmic growth, which is expected to reduce the demand for ribosomes and rRNA. Cells were shifted from rich medium containing dextrose (doubling time of ~90 min) to rich medium containing glycerol (doubling time of ~180 min) and maintained in log phase for 30 doublings. Quantification of *URA3*-marked single-copy ERCs in cells grown in dextrose showed the expected steady-state level of about 20% of the genomic loading control (Figure 3A-B, time point = 0). Upon shift to glycerol, single-copy ERC levels diminished within 7 cell divisions and gradually approached a new steady-state level 2-3 times lower than in the presence of dextrose (Figure 3A-B). The decrease in ERC levels was not the result of repeat loss because the amount of FOA^R^ colonies was indistinguishable before and after growth in glycerol (Figure S3A). The new steady state suggests that ERC levels respond to metabolic demand.

**Figure 3:**
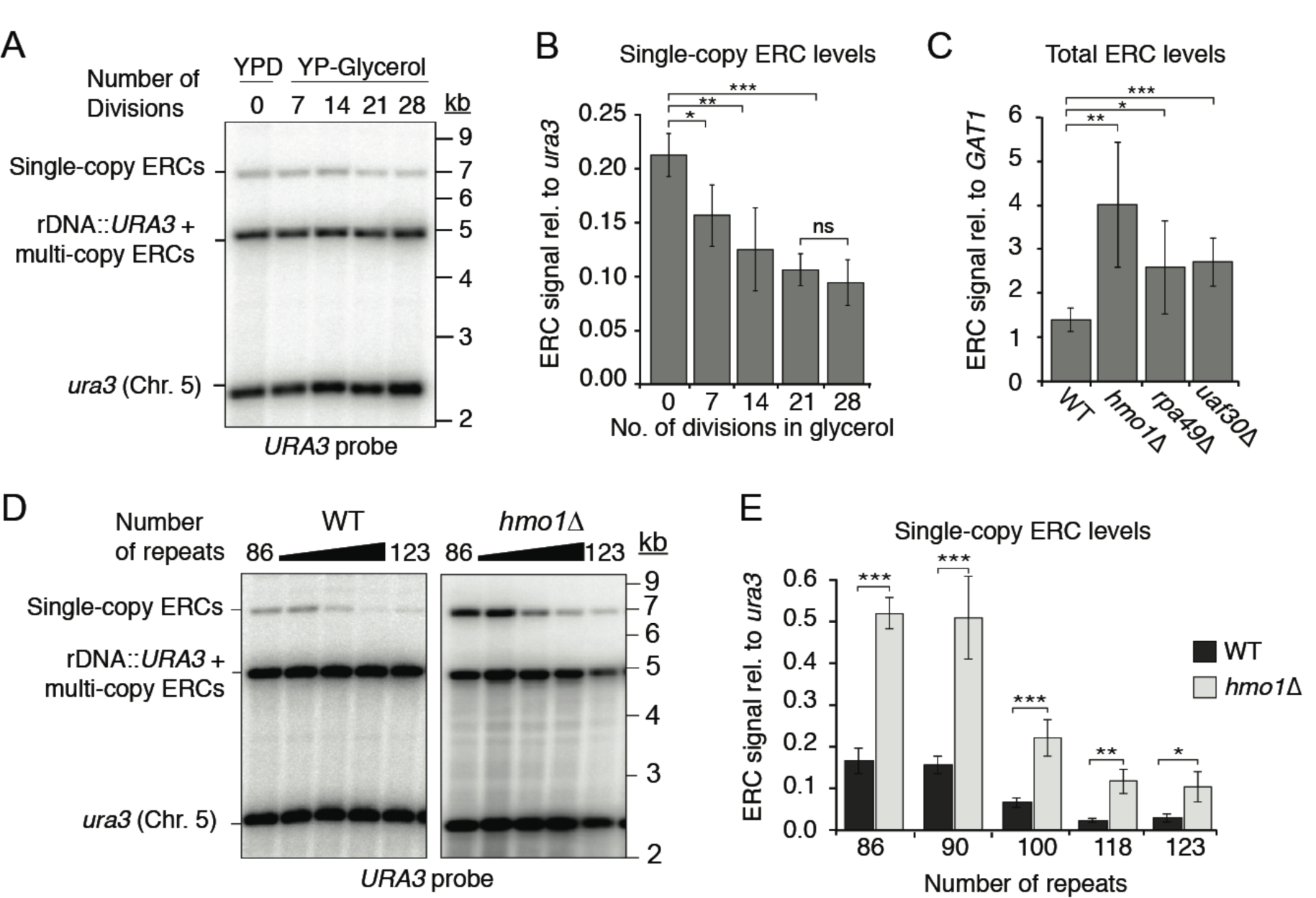
ERC formation is linked to rRNA levels. **(A-B)** Levels *URA3*-marked ERCs (~90-repeat strain) after shift from dextrose (YPD, t=0) to glycerol media. (**A**) Representative Southern blot. (**B**) Average signal of 4 experiments (\**P* < 0.05, \*\**P* < 0.01, \*\*\**P* < 0.001 (paired *t*-test). **(C)** Quantification of ERC levels in ~120-repeat strains (bands a-e; *URA3* probe) relative to *GAT1* upon deletion of *HMO1* (n=4)*, RPA49* (n=6), or *UAF30* (n=4); \**P* < 0.05, \*\**P* < 0.01, \*\*\**P* < 0.001 (two-tailed *t*-test with Holm correction). **(D-E)** ERCs in WT and *hmo1Δ* mutants by arrays size; \*\**P* < 0.01 (two-tailed *t*-test, n=3). WT data in (E) is same as in Figure 1F.

To further test if ERC formation is linked to the demand for rRNA, we reduced rRNA transcription using *hmo1Δ, rpa49Δ,* and *uaf30Δ* mutants, which lack important activators of rRNA transcription (Albert et al., 2013; Gadal et al., 2002; Siddiqi et al., 2001). Under glucose growth conditions, each mutant exhibited a significant increase in steady-state ERC levels (Figures 3C, S3B) despite having array sizes similar to wild type (Figure S3C-D). Analysis of *URA3*-marked repeats similarly revealed a 3-fold increase in steady-state ERC levels in the *hmo1Δ* mutant (Figure 3D-E). Intriguingly, *hmo1Δ* mutants still respond to differences in the size of the chromosomal array, but with a consistent 3-fold overproduction of ERCs. These data suggest that reduced rRNA output leads to miscalibration of the copy-number response. We note that marker loss rate is also increased in *hmo1Δ* mutants (Figure S3E), but this effect is genetically separable. Increased ERC formation in *hmo1Δ* mutants requires *FOB1* (and *RAD52*; Figure S3F), whereas increased marker loss is *FOB1*-independent (Figure S3E). We conclude that ERC production is increased in response to low rRNA levels.

For additional mechanistic insight into the regulation of ERC production, we analyzed binding of Fob1 to the rDNA as a key initial step for ERC production (Figure 4A) (Defossez et al., 1999). Quantitative PCR (qPCR) after chromatin immunoprecipitation (ChIP) revealed that Fob1 binding to the replication-fork barrier (RFB) is significantly increased in short-array strains (Figure 4B), in line with their increased ERC levels. A similar effect is also observed when comparing Fob1 binding to the RFB in short-array strains growing in glycerol versus dextrose (Figure 4C). These data indicate that the observed changes in ERC production are linked to altered Fob1 binding to the RFB.

**Figure 4:**
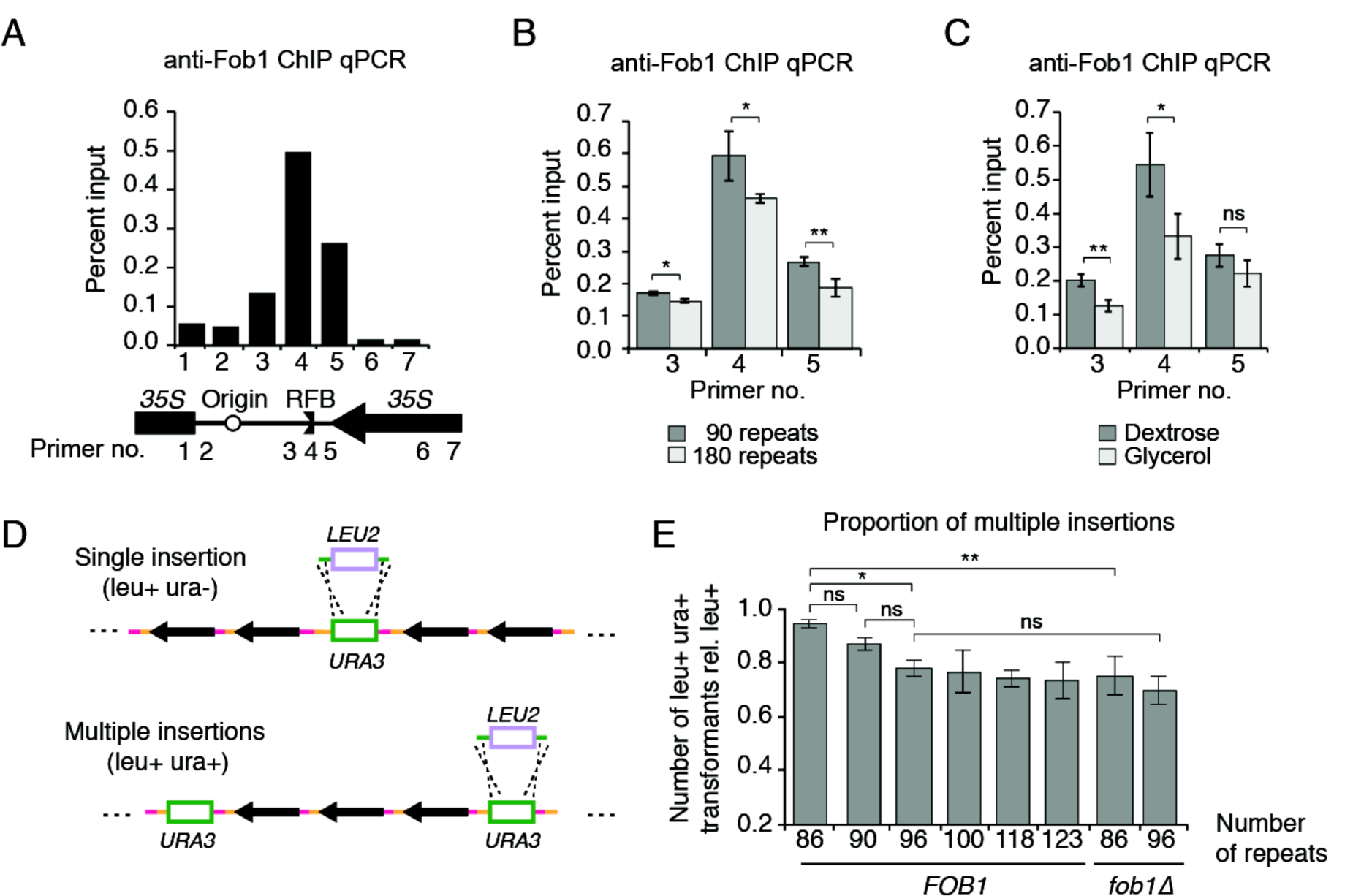
Copy number alters Fob1 binding and ERC reinsertion. **(A-C)** ChIP-qPCR analysis of Fob1 levels in the rDNA **(A)** and at the RFB **(B, C)** in indicated strains and conditions; \**P* < 0.05, \*\**P* < 0.01 (two-tailed *t*-test, n=3). **(D)** Strategy for detecting cells with multiple *URA3* insertions (green) by targeting *URA3* with a *LEU2* construct (blue). **(E)** leu+ ura+ transformants relative to the total leu+ transformants in the indicated strains; \**P* < 0.05, \*\**P* < 0.01 (pair-wise, two-tailed *t*-tests with Holm, n=3).

### Consequences of increased ERC production

Our findings imply a cellular need for compensatory ERC production, even though cells can survive with as little as 20 rDNA repeats (Ide et al., 2010). Strains with very low copy number (20-60) are sensitive to the DNA damaging agent MMS (Ide et al., 2010). However, wild-type and *fob1Δ* strains harboring either 90 or 180 chromosomal rDNA repeats were indistinguishable over a wide range of MMS concentrations (Figure S4A), indicating that ERC formation is dispensable in these situations.

We asked if ERCs might insert into the genome to help re-expand the rDNA array after repeat loss. Such events are difficult to isolate directly because all repeats can serve as potential ERC sources and integration sites, creating substantial heterogeneity. However, reinsertion of a *URA3*-marked ERC should create a duplication of the *URA3* marker elsewhere in the rDNA array that can be captured by targeted *URA3* disruption with *LEU2* (Figure S4B). If multiple *URA3* markers are present, leu+ ura+ transformants will be observed when only one of the *URA3* markers is disrupted (Figure 4D). Applying this assay to six *URA3* insertion strains with rDNA lengths from 86-123 repeats revealed a *FOB1*-dependent, positive correlation between the fraction of leu+ ura+ transformants and ERC levels (Figures 4E, 1E), consistent with dosage-dependent ERC insertion into the rDNA. As marker loss rates were uniform among these strains (Figure 1F), the strain-dependent differences in marker duplication cannot have arisen from unequal sister chromatid exchange. These data suggest that ERCs provide a reservoir for regrowth of shortened rDNA arrays.

## Discussion

Catastrophic loss of repeats by NAHR is an inherent risk to tandem repetitive DNA arrays. Here, we show that *S. cerevisiae* cells respond to large-scale rDNA deletions by proportionally increasing the production of circular rDNA minichromosomes. Intriguingly, although this process also involves NAHR, it does not alter the chromosomal rDNA array. As ERC production relies on DSB formation at the replication fork barrier (Burkhalter and Sogo, 2004), a conceivable model is that the looped-out sequences are re-replicated by the oncoming replication fork (Figure S4C). A similar mechanism has been suggested for the formation of cancer-associated circular EGFR amplicons, which also form without loss of the chromosomal EGFR copies (Vogt et al., 2004). Origin firing in the rDNA, and thus fork-barrier-associated DSB formation, is linked to transcription of the neighboring rRNA gene (Muller et al., 2000), providing a possible explanation for the responsiveness of ERC formation to rRNA transcription.

ERC downregulation likely occurs as a combined consequence of reduced ERC production and passive dilution by cell division. We note that at the cellular level, downregulation is likely limited to newborn daughter cells, as ERCs are almost entirely retained in the mother cell during cell division (Sinclair and Guarente, 1997). This setup allows daughter cells to reassess chromosomal rDNA copy number, but also precludes inheritance of an optimal ERC number, indicating that ERC-mediated copy-number control is non-adaptive.

Extrachromosomal circular DNAs (eccDNAs) are major contributors to tumor heterogeneity (Turner et al., 2017) but are also detected in healthy cells (Shoura et al., 2017). The regulated formation of ERCs indicates that the formation of eccDNAs does not have to be the result of genome instability, providing a framework for understanding the mechanisms driving eccDNA production in healthy cells.

## Acknowledgements

We thank S.P. Bell for the Fob1 antibody, L. Steinmetz for strains, and J. Hubbard, R. Rothstein, M. Siegal, and D. Smith for helpful discussions. This work was supported by NIH grants R01GM088248 and R01GM111715 to A.H. and NIH grant R01CA146940 to H.K. A.R.M. was supported by NIH fellowship T32HD7520-17 and a Dean’s Dissertation Fellowship (NYU).

## Supplementary Materials

**Figure S1**: ERC formation is not linked to the loss of rDNA repeats; related to Figure 1

**Figure S2**: Rapid accumulation of repeat-marked ERCs; related to Figure 2

**Figure S3**: RNAPI mutants increase ERC levels while maintaining array size; related to Figure 3

**Figure S4**: ERC levels are linked to ERC re-insertion frequency; related to Figure 4

**Supplemental Table 1**: Yeast strains used, related to all figures

## Method Details

### Yeast strains, plasmids, transformation, and growth conditions

Strains used in this study are of the SK1 background and are listed in Table S1. To make the collection of strains of different rDNA array sizes, we transformed cells with SphI-digested plasmid pRS306 that had the rDNA intergenic sequence cloned into it between BamHI and EcoRI restriction sites (Fig. 1A-B) (Vader et al., 2011). Lithium acetate transformation was used for insertion of pRS306-rDNA (Fig. 1A-B) and for *URA3*-to-*LEU2* insertion swap (pRS306-to-pRS305) (Fig. 4C-D). Cells were incubated at 30°C in 1M lithium acetate, 50% PEG-3350, 100μg salmon sperm single-stranded DNA, and 200-700 ng of insertion-targeting DNA for 30 minutes prior to a 15 minute heat-shock at 42°C. Due to changes in rDNA size upon lithium acetate transformation (Kwan et al., 2016), electroporation was used for transforming high-copy rDNA plasmid (pRDN-*hyg1 leu2*-D) into a ~90-repeat strain (Fig. 2C-D). Cells were resuspended in chilled 1M sorbitol and subjected to an electro-pulse (1.5kV, 25uF, and 200 Ohms) using a Bio-Rad GenePulser Xcell. When analyzing the effect of plasmid overexpression or mutations, experiments were performed using strains with the same rDNA array size to exclude rDNA size as a variable. All strains were grown at 30°C in 2% YPD, or 3% YPG (glycerol) when indicated.

### Plug preparation and pulsed-field gel electrophoresis conditions

Cultures were grown to saturation and 8.5x10^5^ cells were embedded in agarose using Bio-rad plug molds followed by incubations in zymolyase T100 (200μg/mL) overnight at 37°C, proteinase K (4 mg/mL) overnight at 50°C, and PMSF (1 mM) for one hour at 4°C. PMSF was removed by washing plugs with 1 mL of CHEF TE three times for thirty minutes. Plugs were subsequently run in a 1X TAE, 0.8% agarose gel using a Bio-rad CHEF-DR II at settings: 3V/cm, 50 hrs, 250 seconds start switch time, and 900 seconds end switch time (Figs. 1B, S1A, S1B, S1G, S3C, S3D). *H. wingei* ladder (Bio-rad) was used as size marker.

### Genomic DNA extraction and gel electrophoresis conditions

8.5 x 10^7^ cells undergoing mid-logarithmic growth (OD_600_ 0.5-1.0) were collected and treated with 250 μg/mL zymolyase T100 in 1M sorbitol, 42 mM K_2_HPO_4_, 8mM KH_2_PO_4_, 5mM EDTA at 37°C for 30 minutes, followed by treatment with proteinase K (285 μg) and SDS (0.5%) at 65°C for 1.5 hours. SDS was then precipitated out using potassium acetate. Nucleic acids were ethanol precipitated and treated overnight at 4°C with RNase A (60 μg). DNA was subsequently purified using phenol/chloroform (Roche) and precipitated using isopropanol. Genomic DNA was digested for 4 hours at 37°C using 2μL AfeI (NEB) for Figs. 1C, 2A, S1C, S3B; 4 hours at 37°C with 2μL XhoI (NEB) for Fig. 2C; 4 hours at 37°C with 2μL NdeI + 2μL AvrII (NEB) for Figs. 1E, 3A, 3D, S1H, S2B, S3F; and 24-hours at 37°C with 2μL AfeI + 2μL AvrII + 2μL RecBCD (NEB) for Figs. S1C,D,H. Digested DNA was subsequently run in a 1X TBE, 0.7% agarose gel for 20 hours at 1.5 V/cm.

### Southern blotting, phosphoimaging, and quantification

Southern blotting was performed by alkaline transfer using Hybond-XL membranes (GE). Blots were subsequently probed with ^32^P-labeled DNA complementary to [*GAT1* and *NTS1*] or *URA3.* Signal from blots was detected using FujiFilm imaging plates and simaged using Typhoon FLA9000, at non-saturating conditions for ERC bands and single-copy loading controls. Signals were quantified using ImageJ. Background signal above and below each band was averaged and subtracted from signal of interest to account for well-to-well variation in the amount of DNA shearing. ERC signals were then summed and divided by intensity measurement of single-copy loading controls, *GAT1* or *ura3*.

### Spot tests for growth on 5-FOA and DNA damage sensitivity

Cells were serially diluted 3.5-fold and spotted onto 2% dextrose synthetic complete (SC) media with and without 11.5mM 5-FOA (Toronto Research Chemicals) (Fig. 1G and Fig. S3E). For DNA damage sensitivity assay, cells were similarly spotted onto 2% YPD with and without methyl methanesulfonate (MMS) (Sigma-Aldrich) (Fig. S4).

### Chromatin Immunoprecipitation

8.5 x 10^7^ cells undergoing mid-logarithmic growth (OD_600_ 0.5-1.0) were collected and fixed with 1% formaldehyde for 30 minutes and quenched with glycine. Cells were resuspended in lysis buffer (50mM HEPES/KOH pH 7.5, 140mM NaCl, 1mM EDTA, 1% Triton X-100, 0.1% Na-Deoxycholate, 1mM PMSF, 1mM benzamidine, 1mg/mL bacitracin, Roche Protease Inhibitor tablet) and lysed by glass-bead biopulverizeration (MP Biomedicals). Chromatin was then subjected to shearing using Branson Digital Sonifier Model 250 at 15% amplitude for 15 seconds, repeated five times, and centrifugally separated from cell debris, followed by incubation with Anti-Fob1 (kindly provided by Stephen P. Bell) for 1 hour and a subsequent incubation with magnetic protein G beads (NEB) for 1 hour. Immunoprecipitated chromatin was then washed with lysis buffer, high salt buffer, lithium chloride buffer, and TE, followed by elution at 65°C and DNA purification using Qiagen PCR purification columns. Quantitative PCR was performed on both immunoprecipitated and non-immunoprecipitated input samples, and then quantified using percent input method: 100^*^2^(Adjusted input – IP).

### Assay for detection of multiple rDNA insertions

Cells containing an integrated *URA3* marker were transformed with pRS305 that had been digested at BamHI (NEB) and HindIII (NEB) sites. Positive transformants were grown on leucine dropout media, and after three days of growth at 30°C, selection plates were replica-plated onto uracil dropout media and 11.5mM 5-FOA. The proportion of leu+ ura+ transformations was quantified relative to the total number of leu+ transformants. A total of 450 or more leu+ transformants were scored for each strain from three transformations. Statistical comparisons were performed using two-tailed, pairwise *t*-tests with a Holm correction for multiple testing.

### rDNA expansion experiment

Strains were bottlenecked by streaking for single colonies on 2% YPD and incubated for 2 days. The number of vegetative cell divisions were estimated by colony size (2mm was equivalent to approximately 20 cell divisions) (*7*).

### RNA preparation and qPCR analysis

10μg of total RNA was phenol/chloroform extracted and treated with TURBO DNase (Invitrogen, 1μL per sample) at 37°C for 1 hour. DNase inactivation reagent (Invitrogen, 5μL per sample) was added and incubated with samples for 5 minutes at room temperature. 1μg of this RNA was used for reverse transcription (RT). Samples for RT were incubated with Revert Aid (Fermentas, 1μL per sample), Revert Aid buffer (Fisher), Oligo-dT (100μM, Fisher, 1μL per sample) and RNaseOut recombinant ribonuclease inhibitor (Invitrogen, 1μL per sample) at 42°C for 1 hour in a 20μL reaction. Samples without RT were run to control for the DNase step. Reactions were terminated by incubation at 70°C for 10 minutes. Samples were diluted to 400μL for analysis by qPCR. For qPCR analysis, 8μL samples (diluted ChIP samples or diluted RT samples) and 1μL of each primer (20nM) were mixed with Quantifast SYBR green master mix (Qiagen, 10μL per reaction) and placed in a BioRad CFX96 Real-Time System. Program: 95°C, 5 minutes; 39 cycles: 95°C, 10 seconds; 60°C, 30 seconds.

### Statistical analyses

For analyses of both total ERC levels and *URA3*-marked ERCs as a function of rDNA array size, *P* < 4.5x10^-5^ was determined by Spearman’s rank correlation tests. These data also showed high correlations to power law functions as determined by coefficient of determination (r^2^) values. Error bars represent standard deviations of measurements in all main and supplemental figures.

**Figure S1:**
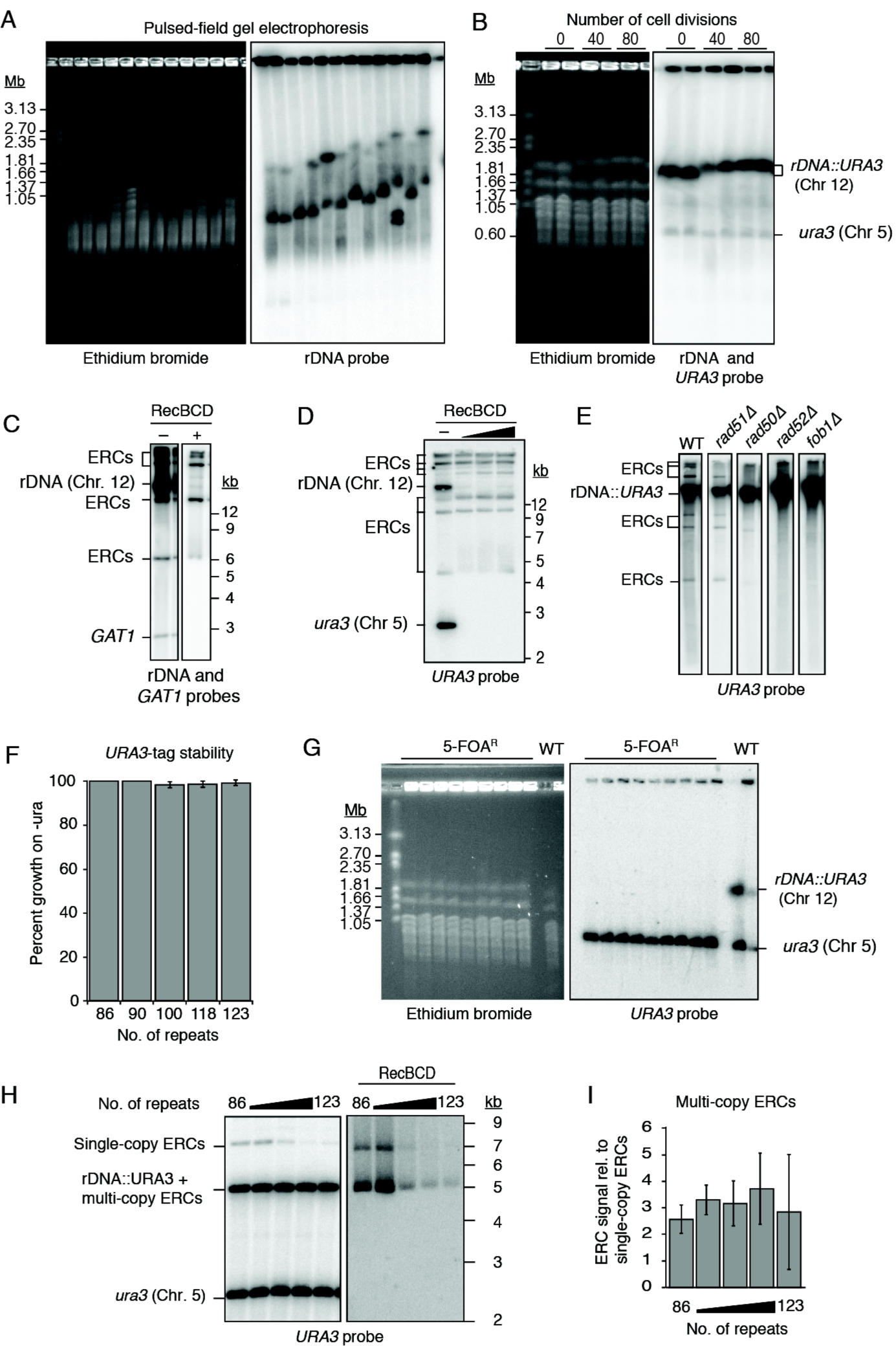
ERC formation is not linked to the loss of rDNA repeats. (A) PFGE of *URA3*-marked strains digested with AfeI. Lanes ordered roughly by rDNA array repeat no. from left to right (93, 89, 96, 100, 123, 118, 145, 133, 153, 179, 147, 183). (B) PFGE of ~90-repeat strain after 40 and 80 cell divisions on YPD. (C-E) Southern analyses of ERCs (F) Proportion of *URA3*-marked cells that grow on uracil-depleted meida (n=250 for each strain). (G) PFGE of 5-FOA^R^ strains derived from ~90-repeat *URA3*-marked strain; n=9. (H-I) Southern analysis of multi-copy ERCs by RecBCD exonuclease treatment; n=3.

**Figure S2:**
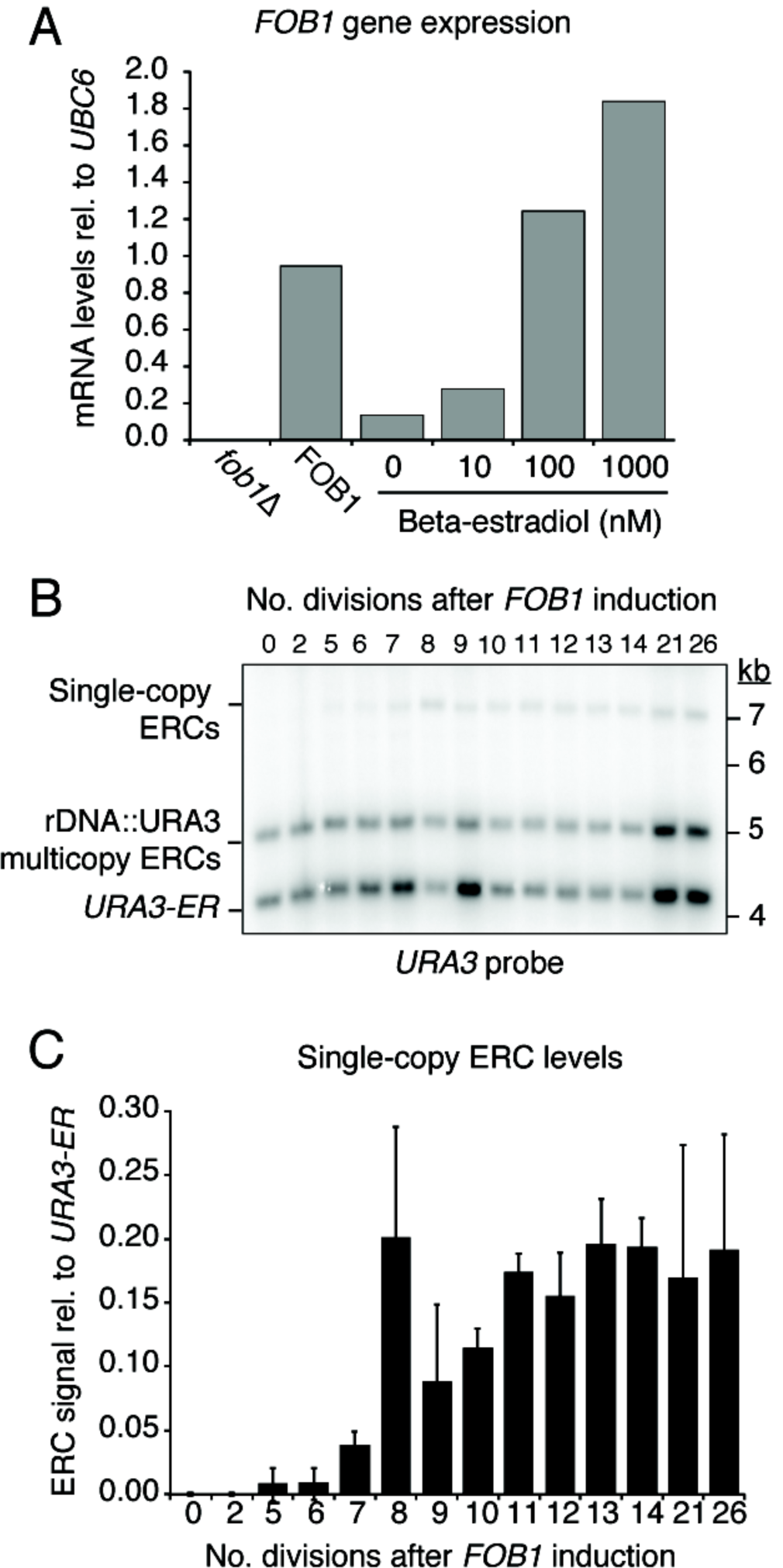
Rapid accumulation of repeat-marked ERCs. (A) Quantification of *FOB1* mRNA levels by qPCR of strains with *fob1*Δ mutation, WT *FOB1* gene, and the Gal-inducible *FOB1* strain; n=2. The latter strain was treated with varying concentrations of Beta-estradiol, whereas the former two were subject to mock treatment with ethanol. Each strain tested contains ~90-repeats. (B-C) Detection and quantification of single-copy ERC kinetics in a ~90-repeat strain over the course of 26 cell divisions before (t=0) and after treatment with 100nM Beta-estradiol by Southern blotting and *URA3* probing; n=3.

**Figure S3:**
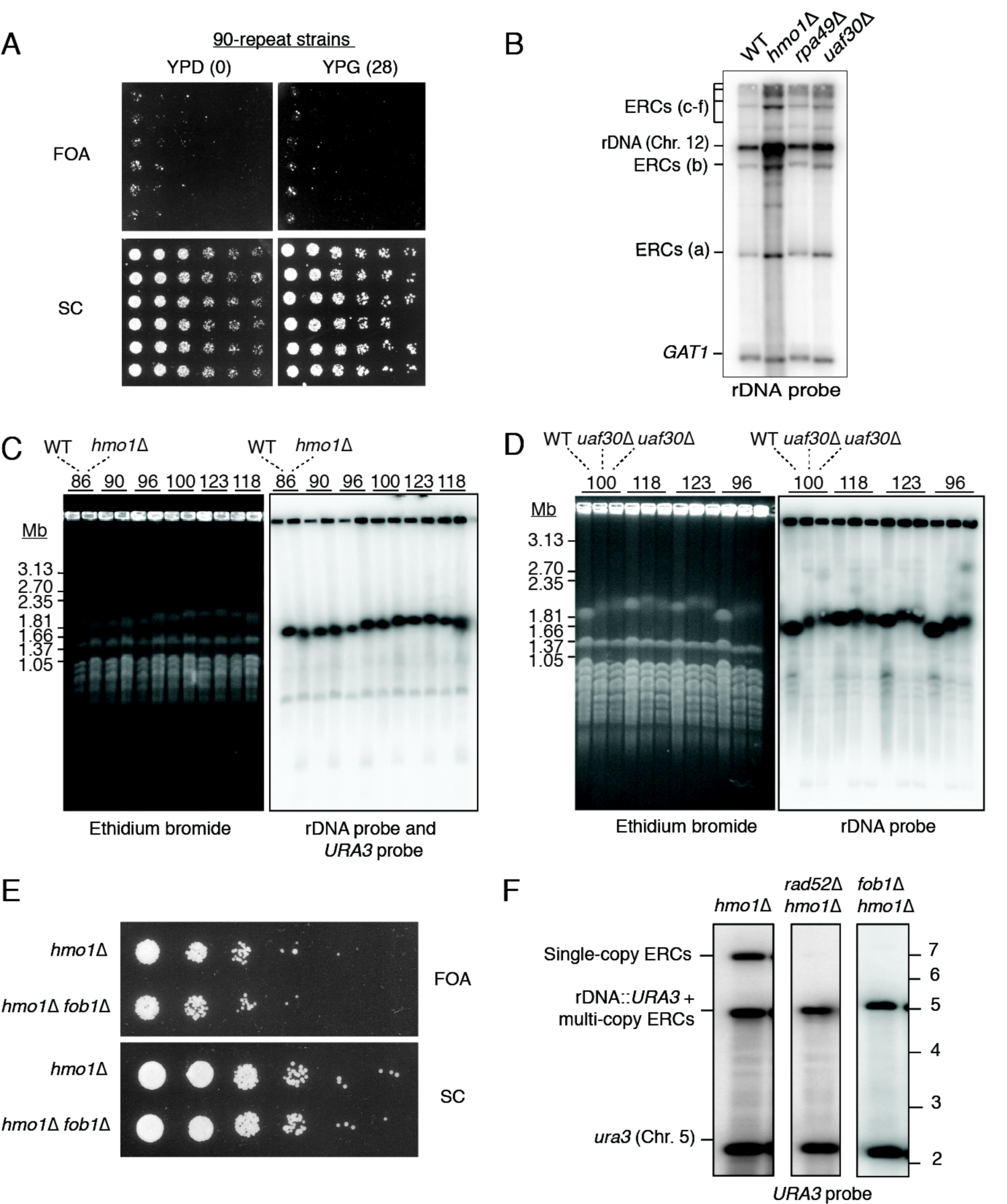
RNAPI mutants increase ERC levels while maintaining array size. (A) Spot assay of ~90-repeat, *URA3*-marked strains on synthetic complete (SC) media with and without 5-FOA (3.5-fold dilutions) before (YPD) and after 28 cell divisions on glycerol (YPG). (B) Detection of total ERCs in WT and RNAPI mutants *hmo1*Δ and *uaf30*Δ by Southern blotting and probing with rDNA sequence. (C-D) PFGE of *hmo1*Δ and *uaf30*Δ strains used in Fig. 3C and Fig. S3B; probed against rDNA and *URA3* sequences. Samples were grouped by repeat number with WT followed by *hmo1*Δ samples or WT followed by two *uaf30*Δ clones. (E) Spot assay of *URA3*-marked *hmo1*Δ and *hmo1*Δ *fob1*Δ strains on synthetic complete (SC) media with and without 5-FOA (3.5-fold dilutions). (F) Southern analysis of *URA3*-marked ERCs in *hmo1*Δ, *hmo1*Δ *fob1*Δ, and *hmo1*Δ *rad52*Δ.

**Figure S4:**
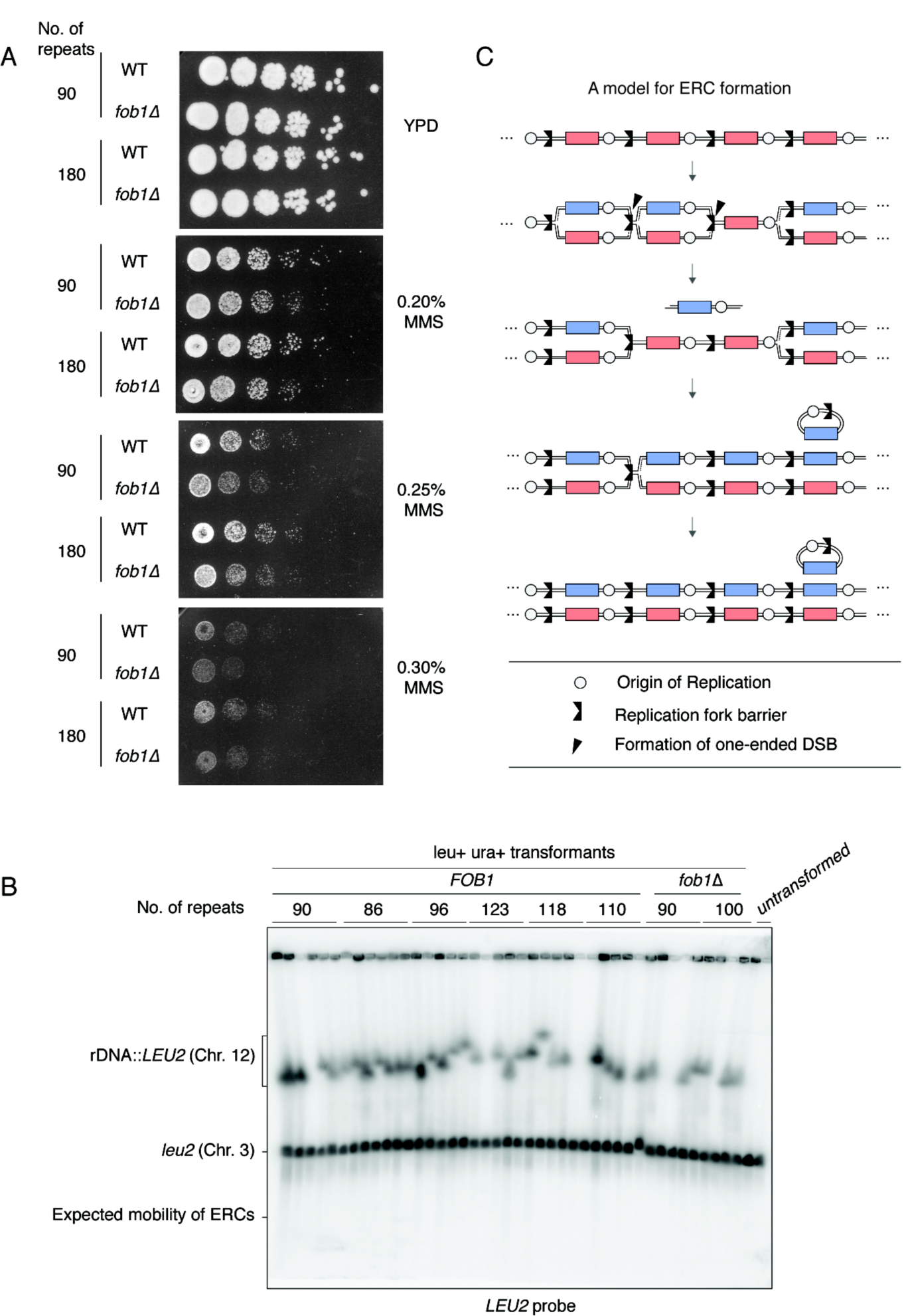
ERC levels are linked to ERC re-insertion frequency. **(A)** Spot assay of 90‐ and 180-repeat *URA3*-marked strains on rich media containing varying amounts of MMS (5-fold dilutions). **(B)** Confirmation of integration of *LEU2* construct at the rDNA by PFGE of undigested plugs and probing for *LEU2* sequence. **(C)** A model for ERC formation that was adapted (Vogt et al., 2004) and modified to include the unipolar directionality of rDNA replication (Brewer and Fangman, 1988).

**Table S1.**

| Strain | Genotype | rDNA array repeat no. | Used in Figure Panel |
| --- | --- | --- | --- |
| H5975 | MATa, lys2, ho::LYS2, trp1::hisG, his3::hisG, leu2::hisG, ura3, rDNA::URA3 | 86 | 1B-G, 3D-E, 4A, 4C, S1A, S1F, S1H-I, S3C-D |
| H7592 | MATa, lys2, ho::LYS2, trp1::hisG, his3::hisG, leu2::hisG, ura3, rDNA::URA3 | 90 | 1B-G, 3A-B, 3D-E, 4A-C, S1A-B, S1D-I, S2A, S3A, S3C-D |
| H6375 | MATa, lys2, ho::LYS2, trp1::hisG, his3::hisG, leu2::hisG, ura3, rDNA::URA3 | 96 | 1B-G, 3D-E, S1A, S1F, S1H-I, S3C-D |
| H5979 | MATa, lys2, ho::LYS2, trp1::hisG, his3::hisG, leu2::hisG, ura3, rDNA::URA3 | 100 | 1B-G, 3D-E, 4A-C, S1A, S1F, S1H-I, S3C-D |
| H6373 | MATa, lys2, ho::LYS2, trp1::hisG, his3::hisG, leu2::hisG, ura3, rDNA::URA3 | 118 | 1B-G, 3C, 3D-E, S1A, S1F, S1H-I, S3B-D |
| H6374 | MATalpha, lys2, ho::LYS2, trp1::hisG, his3::hisG, leu2::hisG, ura3, rDNA::URA3 | 123 | 1B-G, 3C, 3D-E, S1A, S1F, S1H-I, S3B-D |
| H9309 | MATalpha, lys2, ho::LYS2, trp1::hisG, his3::hisG, leu2::hisG, ura3, rDNA::URA3 | 133 | 1B-D, S1A |
| H9311 | MATalpha, lys2, ho::LYS2, trp1::hisG, his3::hisG, leu2::hisG, ura3, rDNA::URA3 | 145 | 1B-D, S1A |
| H9310 | MATalpha, lys2, ho::LYS2, trp1::hisG, his3::hisG, leu2::hisG, ura3, rDNA::URA3 | 147 | 1B-D, S1A |
| H9312 | MATalpha, lys2, ho::LYS2, trp1::hisG, his3::hisG, leu2::hisG, ura3, rDNA::URA3 | 153 | 1B-D, S1A |
| H9313 | MATalpha, lys2, ho::LYS2, trp1::hisG, his3::hisG, leu2::hisG, ura3, rDNA::URA3 | 179 | 1B-D, 4C, S1A |
| H9314 | MATalpha, lys2, ho::LYS2, trp1::hisG, his3::hisG, leu2::hisG, ura3, rDNA::URA3 | 183 | 1B-D, 4C, S1A |
| H8535 | MATa, ho::LYS2, lys2, ura3, leu2::hisG, his3::hisG, trp1::hisG, ura3::pGPD1-GAL4(848).ER::URA3, HisMX::pGAL1::Fob1; flo8Δ(unmarked), rDNA::URA3 | 90 | 2A-B, S2A-B |
| H9296 | MATa, lys2, ho::LYS2, trp1::hisG, his3::hisG, leu2::hisG, ura3 | 90 | 2C-D, S1G |
| H9307 | MATa, lys2, ho::LYS2, trp1::hisG, his3::hisG, leu2::hisG, ura3, [pRDN-hyg::URA3::leu2-8] | 90 | 2C-D |
| H8166 | MATa, lys2, ho::LYS2, trp1::hisG, his3::hisG, leu2::hisG, ura3, rDNA::URA3, rDNA::URA3, hmo1Δ::KanMX4, rDNA::URA3, hmo1Δ::KanMX4 | 86 | 3D-E, S3C-D, S3F |
| H8167 | MATa, lys2, ho::LYS2, trp1::hisG, his3::hisG, leu2::hisG, ura3, rDNA::URA3, rDNA::URA3, hmo1Δ::KanMX4, rDNA::URA3, hmo1Δ::KanMX4 | 90 | 3D-E, S3C-D |
| H8168 | MATa, lys2, ho::LYS2, trp1::hisG, his3::hisG, leu2::hisG, ura3, rDNA::URA3, rDNA::URA3, hmo1Δ::KanMX4, rDNA::URA3, hmo1Δ::KanMX4 | 96 | 3D-E, S3C-D |
| H8169 | MATa, lys2, ho::LYS2, trp1::hisG, his3::hisG, leu2::hisG, ura3, rDNA::URA3, rDNA::URA3, hmo1Δ::KanMX4, rDNA::URA3, hmo1Δ::KanMX4 | 118 | 3C-E, S3B-D |
| H8170 | MATa, lys2, ho::LYS2, trp1::hisG, his3::hisG, leu2::hisG, ura3, rDNA::URA3, rDNA::URA3, hmo1Δ::KanMX4, rDNA::URA3, hmo1Δ::KanMX4 | 123 | 3C-E, S3B-D |
| H9425 | MATa, lys2, ho::LYS2, trp1::hisG, his3::hisG, leu2::hisG, ura3, rDNA::URA3, uaf30D::KanMX | 118 | 3C, S3B-D |
| H9426 | MATa, lys2, ho::LYS2, trp1::hisG, his3::hisG, leu2::hisG, ura3, rDNA::URA3, uaf30D::KanMX | 123 | 3C, S3B-D |
| H9427 | MATa, lys2, ho::LYS2, trp1::hisG, his3::hisG, leu2::hisG, ura3, rDNA::URA3, uaf30D::KanMX | 125 | 3C, S3B-D |
| H8226 | MATa, lys2, ho::LYS2, trp1::hisG, his3::hisG, leu2::hisG, ura3, rDNA::URA3, flo8Δ (unmarked) | 86 | 4D, S4A |
| H8227 | MATa, lys2, ho::LYS2, trp1::hisG, his3::hisG, leu2::hisG, ura3, rDNA::URA3, flo8Δ (unmarked) | 90 | 4D |
| H8228 | MATa, lys2, ho::LYS2, trp1::hisG, his3::hisG, leu2::hisG, ura3, rDNA::URA3, flo8Δ (unmarked) | 96 | 4D |
| H8229 | MATa, lys2, ho::LYS2, trp1::hisG, his3::hisG, leu2::hisG, ura3, rDNA::URA3, flo8Δ (unmarked) | 100 | 4D |
| H8230 | MATa, lys2, ho::LYS2, trp1::hisG, his3::hisG, leu2::hisG, ura3, rDNA::URA3, flo8Δ (unmarked) | 118 | 4D |
| H8231 | MATa, lys2, ho::LYS2, trp1::hisG, his3::hisG, leu2::hisG, ura3, rDNA::URA3, flo8Δ (unmarked) | 123 | 4D |
| H7745 | MATa, lys2, ho::LYS2, trp1::hisG, his3::hisG, leu2::hisG, ura3, rDNA::URA3, flo8Δ (unmarked), fob1Δ::KanMX | 86 | 4D, S1E, S2A, S4A |
| H7749 | MATa, lys2, ho::LYS2, trp1::hisG, his3::hisG, leu2::hisG, ura3, rDNA::URA3, flo8Δ (unmarked), fob1Δ::KanMX | 96 | 4D |
| H8253 | MATa, lys2, ho::LYS2, trp1::hisG, his3::hisG, leu2::hisG, ura3, rDNA::URA3, rad51Δ | 90 | S1E |
| H9317 | MATa, lys2, ho::LYS2, trp1::hisG, his3::hisG, leu2::hisG, ura3, rDNA::URA3, rad50Δ, flo8Δ (unmarked) | 90 | S1E |
| H8204 | MATa, lys2, ho::LYS2, trp1::hisG, his3::hisG, leu2::hisG, ura3, rDNA::URA3, rad52Δ | 90 | S1E |
| H6703 | MATa, lys2, ho::LYS2, trp1::hisG, his3::hisG, leu2::hisG, ura3, rDNA::URA3, fob1Δ::KanMX | 96 | S3E |
| H8683 | MATa, lys2, ho::LYS2, trp1::hisG, his3::hisG, leu2::hisG, ura3, rDNA::URA3, fob1Δ::KanMX, hmo1Δ::KanMX4 | 90 | S3E |
| H8687 | MATa, lys2, ho::LYS2, trp1::hisG, his3::hisG, leu2::hisG, ura3, rDNA::URA3, fob1Δ::KanMX, hmo1Δ::KanMX4 | 96 | S3E |
| H6690 | MATa, lys2, ho::LYS2, trp1::hisG, his3::hisG, leu2::hisG, ura3, rDNA::URA3, fob1Δ::KanMX | 90 | S3E |
| H8242 | MATa, lys2, ho::LYS2, trp1::hisG, his3::hisG, leu2::hisG, ura3, rDNA::URA3, hmo1Δ::KanMX, rad52Δ::TRP1 | 90 | S3F |
| H9328 | MATalpha, lys2, ho::LYS2, trp1::hisG, his3::hisG, leu2::hisG, ura3, rDNA::URA3 | 183 | S4A |
| H8796 | MATalpha, lys2, ho::LYS2, trp1::hisG, his3::hisG, leu2::hisG, ura3, rDNA::URA3, fob1Δ::KanMX | 183 | S4A |

